# The chloroplastic phosphate transporter CrPHT4-7 supports phosphate homeostasis and photosynthesis in Chlamydomonas

**DOI:** 10.1101/2023.09.08.556869

**Authors:** Dávid Tóth, Soujanya Kuntam, Áron Ferenczi, André Vidal-Meireles, László Kovács, Lianyong Wang, Zsuzsa Sarkadi, Ede Migh, Klára Szentmihályi, Roland Tengölics, Juliane Neupert, Ralph Bock, Martin C. Jonikas, Attila Molnar, Szilvia Z. Tóth

## Abstract

In eukaryotic cells, phosphorus is assimilated and utilized primarily as phosphate (Pi). Pi homeostasis is mediated by transporters that have not yet been adequately characterized in green algae. This study reports on CrPHT4-7 from *Chlamydomonas reinhardtii*, a member of the PHT4 transporter family, which exhibits remarkable similarity to AtPHT4;4 from *Arabidopsis thaliana*, a chloroplastic ascorbate transporter. Using fluorescent protein tagging we show that CrPHT4-7 resides in the chloroplast envelope membrane. *Crpht4-7* mutants, generated by the CRISPR/Cas12a-mediated single-strand templated repair, show retarded growth especially in high light, enhanced sensitivity to phosphorus limitation, reduced ATP level, strong ascorbate accumulation and diminished non-photochemical quenching in high light. Conversely, CrPHT4-7 overexpressing lines exhibit enhanced biomass accumulation under high light conditions in comparison with the wild-type strain. Expressing CrPHT4-7 in a yeast strain lacking Pi transporters substantially recovered its slow growth phenotype demonstrating that it transports Pi. Even though CrPHT4-7 shows a high degree of similarity to AtPHT4;4, it does not display any significant ascorbate transport activity in yeast or intact algal cells. Thus, the results demonstrate that CrPHT4-7 functions as a chloroplastic Pi transporter essential for maintaining Pi homeostasis and photosynthesis in *Chlamydomonas reinhardtii*.

**One-sentence summary:** We demonstrate that the CrPHT4-7 transporter of *Chlamydomonas reinhardtii* is located in the chloroplast envelope membrane and contributes to maintaining phosphate homeostasis and photosynthesis.

## Introduction

Phosphorus is essential for living organisms and is found in every compartment of the plant cell. It is a structural component of nucleic acids and phospholipids, and also indispensable for signal transduction and energy transfer reactions, including photosynthesis. Plants take up phosphorus from the soil in the form of inorganic phosphate (Pi) through the cell wall and plasma membrane, which then is transported into the various cell organelles. Despite its widespread occurrence in the environment, Pi availability often limits plant growth, because of phosphate complexation with metal cations and organic particles in the soil (e.g., Gutiérrez-Alanís et al., 2018, Crombez et al., 2019). Fertilizers, derived from non-renewable rock phosphate, improve crop yields that otherwise are limited by Pi availability, but the leaching of excess Pi into aquatic ecosystems causes environmental problems such as eutrophication. For these reasons, studying Pi uptake and transport in plants is of high importance.

PHT family members are the best-studied phosphate transporters in vascular plants. They are well known for their roles in Pi uptake from soil and Pi translocation within the plant (Versaw and Garcia, 2017; Wang et al., 2021). *Arabidopsis thaliana* has five high-affinity Pi transporter families (PHT1-5) that are distinguished based on their functional differences and subcellular localization. The PHT1 proteins are plasma membrane proton-coupled Pi-symporters that mediate Pi acquisition from the soil and Pi translocation within the plant. Members of the PHT2 and PHT4 families are present in plastids and in the Golgi apparatus, whereas PHT3 transporters are found in mitochondria and PHT5;1 is a vacuolar Pi transporter (Versaw and Garcia, 2017; Srivastava et al., 2018).

Phosphate transport is poorly studied in green algae and surprisingly, no Pi transporter has been characterized in detail (Wang et al., 2020). Understanding the mechanisms of Pi uptake and cellular distribution is highly relevant since microalgae can accumulate and store large amounts of Pi in the form of polyphosphate granules in specific vacuoles called acidocalcisomes (Sanz-Luque et al. 2020). This so-called ‘‘luxury uptake’’ (Riegman et al., 2000) may enable recovery of Pi upon wastewater treatment (Shilton et al., 2012) to subsequently produce phosphate-rich fertilizers (Slocombe et al., 2020). Thus, understanding Pi uptake and transport in microalgae are of high importance to the protection of the environment and water management.

The *PHT* gene family in *C. reinhardtii* contains 25 putative *PHT* genes, categorized in four subfamilies, namely *CrPTA*, *CrPTB*, *CrPHT3*, and *CrPHT4* (Wang et al., 2020). The *CrPTA, CrPTB*, *CrPHT3*, and *CrPHT4* subfamilies may contain four, eleven, one, and nine members, respectively (Wang et al 2020). Members of the CrPTA family, a sister family of PHT1 in land plants, may be found in the plasma membrane (Wang et al., 2020) or targeted to secretory and other pathways (Tardif et al., 2012, Wang et al., 2023). CrPTB members were shown or predicted to be targeted to the secretory and other pathways (Tardif et al., 2012, Wang et al., 2020, Wang et al., 2023); however, based on homology with streptophyte algae, they are likely to be located in the plasma membrane (Bonnot et al., 2017). CrPHT3 (Cre03.g172300) is possibly found in mitochondria (Tardif et al., 2012, Wang et al., 2020, Wang et al., 2023). Several CrPHT4 family members are predicted to be localized in the chloroplast, whereas others may be targeted to secretory pathways or the mitochondria (Tardif et al., 2012, Wang et al., 2020, Wang et al., 2023). It is interesting to note that CrPHT transcript levels responded differently to Pi starvation, with most genes belonging to the *CrPTA* and *CrPTB* families showing significant inductions (Moseley et al., 2006, Wang et al., 2020).

Here, we investigated a member of the CrPHT4 family, called CrPHT4-7 (Cre16.g663600, called CrPHT7 in Phytozome v. 13). This transporter has several PHT4 homologs in *Arabidopsis thaliana* with varied location and roles: *AtPHT4;1* to *AtPHT4;5* are expressed in plastids, whereas *AtPHT4;6* in the Golgi apparatus (Guo et al., 2008a; reviewed by Versaw and Garcia, 2017, Fabiańska et al., 2019). AtPHT4;1 was found in the thylakoid membranes (Pavón et al., 2008), and AtPHT4;4 in the chloroplast envelope membrane of mesophyll cells (Miyaji et al., 2015). The expressions of *AtPHT4;3* and *AtPHT4;5* are restricted mostly to leaf phloem cells, and *AtPHT4;2* is most highly expressed in the roots and other non-photosynthetic tissues (Guo et al., 2008b). All AtPHT4 transporters may act as phosphate transporter as they could complement the yeast PAM2 mutant lacking Pi transporters (Guo et al., 2008a), and they exhibit H^+^ and/or Na^+^-coupled Pi transport activities (Guo et al., 2008a, Irigoyen et al., 2011, Miyaji et al., 2015). Interestingly, it was found that AtPHT4;4 transports ascorbate (Asc) into the chloroplasts (Miyaji et al., 2015), to ensure appropriate Asc level for its multiple roles (Tóth, 2023). AtPHT4;1, on the other hand, may export Pi out of the thylakoid lumen (Karlsson et al., 2015). AtPHT4;2 has been shown to act bidirectionally, and its suggested physiological role is to export Pi from root plastids to support ATP homeostasis (Irigoyen et al., 2011).

Here we found that CrPHT4-7 is a Pi transporter located in the chloroplast envelope membrane of *C. reinhardtii*, and it is required for maintaining Pi homeostasis and optimal photosynthesis under high light conditions.

## Results

### CrPHT4-7 is localized in the chloroplast envelope membrane

CrPHT4-7 belongs to the PHT4 family of transporters, showing similarity to members of the solute carrier family 17 (sodium-dependent Pi co-transporter, SLC17A). CrPHT4-7 shows 42,6% similarity to the *Arabidopsis thaliana* AtPHT4;5 (AT5G20380) Pi transporter and around 29 to 36% similarity with other Pi transporters in the PHT4 family, namely AtPHT4;2, 4;1, 4;6, and 4;3. CrPHT4-7 also shows a relatively high, 37,4% similarity to the chloroplastic Asc transporter AtPHT4;4 (AT4G00370.1) (according to Phytozome v.13, see Suppl. Fig. 1 for the sequence alignments).

In *Arabidopsis*, AtPHT4 transporters are located in the chloroplast envelope membrane of plastids, in thylakoid membranes, and in the Golgi apparatus (reviewed by Fabiańska et al., 2019). Prediction algorithms do not provide a clear indication as to where CrPHT4-7 is localized within the cell. According to DeepLoc 1.0 (Thumuluri et al., 2022), CrPHT4-7 is associated with the Golgi apparatus, whereas LocTree 3 (Goldberg et al., 2014) predicts that the mature protein is localized in the chloroplast membrane. In contrast, ChloroP 1.1 (Emanuelsson et al., 1999) indicates that CrPHT4-7 is not targeted to the chloroplast, and PredAlgo 1.0 (Tardif et al., 2012) predicts that is not in the chloroplast, mitochondria or secretory pathway. The *in silico* analysis by Wang et al. (2020) suggested that CrPHT4-7 is likely localized in the secretory pathway. The recently developed protein prediction tool PB-Chlamy predicts that PHT4-7 is found in the chloroplast (Wang et al., 2023).

To determine the subcellular location of CrPHT4-7, we tagged the CrPHT4-7 with the fluorescent marker Venus at the C-terminus and then introduced the resulting construct (pLM005-CrPHT4-7, Fig. 1A) into the UVM11 strain that has been shown to support enhanced transgene expression (Neupert et al., 2009, Neupert et al., 2020). With its faster maturation rate, improved folding, and reduced sensitivity to environmental pH, Venus represents a versatile fluorescent protein tag (Nagai et al., 2002; the pLM005 base plasmid has been employed in Wang et al., 2023). In parallel, we also introduced the construct into the *Chlamydomonas* CC-4533 strain (also called cMJ030), which is the host strain used in the Chlamydomonas Library Project (Fauser et al., 2022, Wang et al., 2023). In both the UVM11 and CC-4533 strains, the fluorescent signals from Venus-tagged CrPHT4-7 could be detected (Fig. 1, and Suppl. Fig. 2, respectively). In the case of the UVM11 strain, the signal was detected in 41 out of 93 transformed clones tested (corresponding to 44% efficiency). The merged images of Venus-tagged CrPHT4-7 and Chl *a* autofluorescence (Fig. 1, Suppl. Fig. 2) show that CrPHT4-7 is localized to the chloroplast envelope.

**Figure 1.**
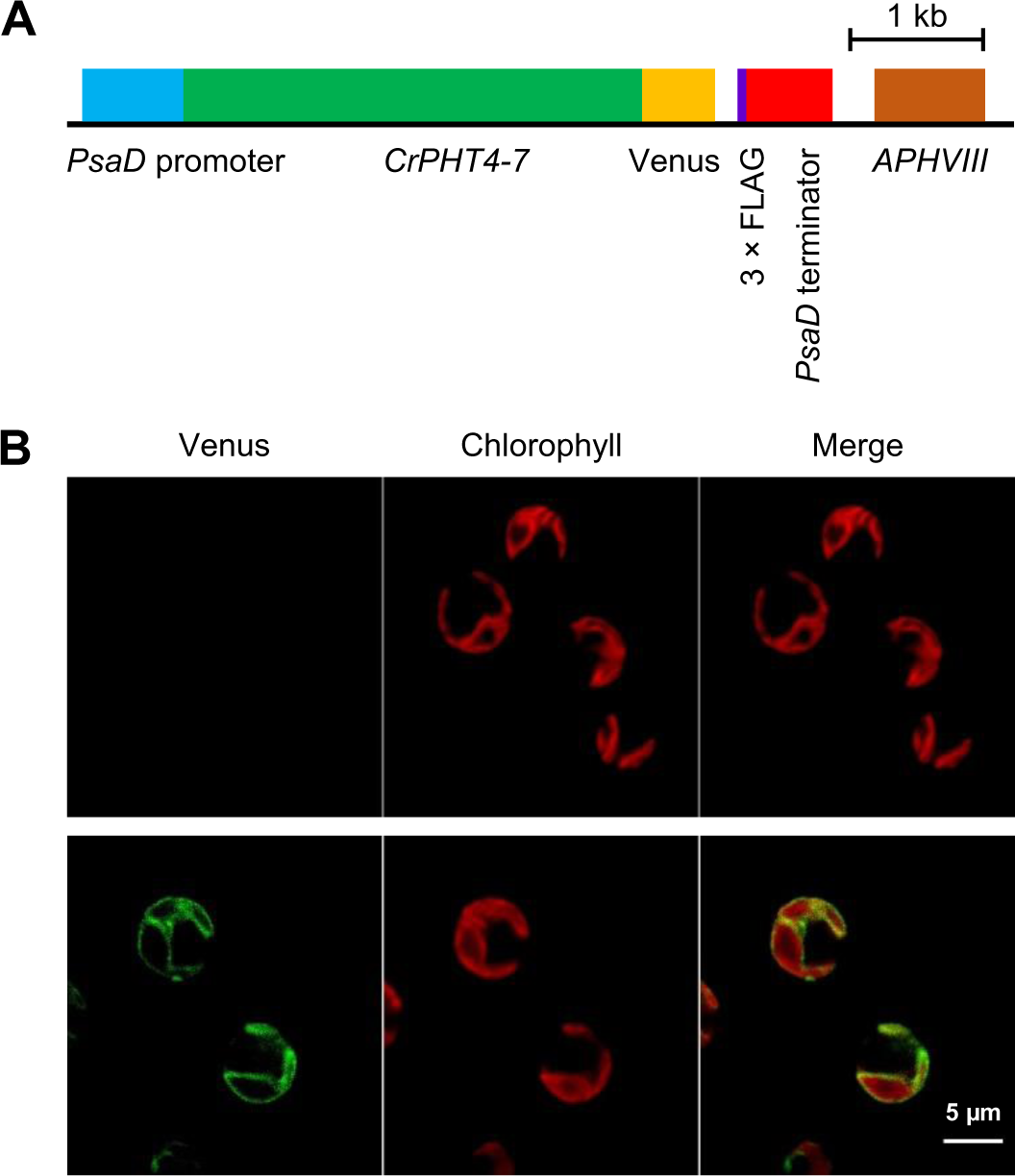
CrPHT4-7 is found in the chloroplast envelope membrane. **A,** Map of the pLM005-CrPHT4-7 plasmid expressing a Venus-tagged CrPHT4-7 version. **B,** Representative fluorescence microscopic images of the UVM11 strain (upper row) and the UVM11 strain expressing pLM005-CrPHT4-7 with Venus-3×FLAG (lower row). Venus fluorescence and Chl auto-fluorescence were detected between 520-540 nm and 650-750 nm, respectively. The merged Venus + Chl fluorescence image is also shown. Scale bar: 5 μm.

### CrPHT4-7 is required for normal growth especially at high light

To investigate the physiological role of CrPHT4-7, we studied *pht4-7* knock out mutants, which were generated by CRISPR/Cas12a-mediated single-strand templated repair introducing early stop codons (Ferenczi et al., 2017). In the initial CRISPR/Cas12a-ssODN mutagenesis screen, the *pht4-7* mutants formed smaller colonies than wild-type cells (WT, CC-1883); Ferenczi et al., 2017). In agreement with this observation, five independent mutant lines showed a similar slow growth phenotype in comparison with the WT strain as estimated by absorbance at 720 nm (OD_720_) in a Multi-Cultivator photobioreactor (Suppl. Fig. 3; Thoré et al., 2021). Of these, we have randomly selected two independent mutants, called *pht4-7#7* and *#9* for further detailed analyses.

The presence of the introduced sequence variations and premature stop codons was confirmed by Sanger sequencing in the *pht4-7#7* and *#9* mutant lines (Fig. 2A). The stop codons were introduced into the third exon of *CrPHT4-7* to prevent the translation of half of the C-terminal transmembrane helices (Fig. 2B). Consequently, the *pht4-7#7* and *#9* mutants are very likely to express a strongly truncated, non-functional form of CrPHT4-7.

**Figure 2.**
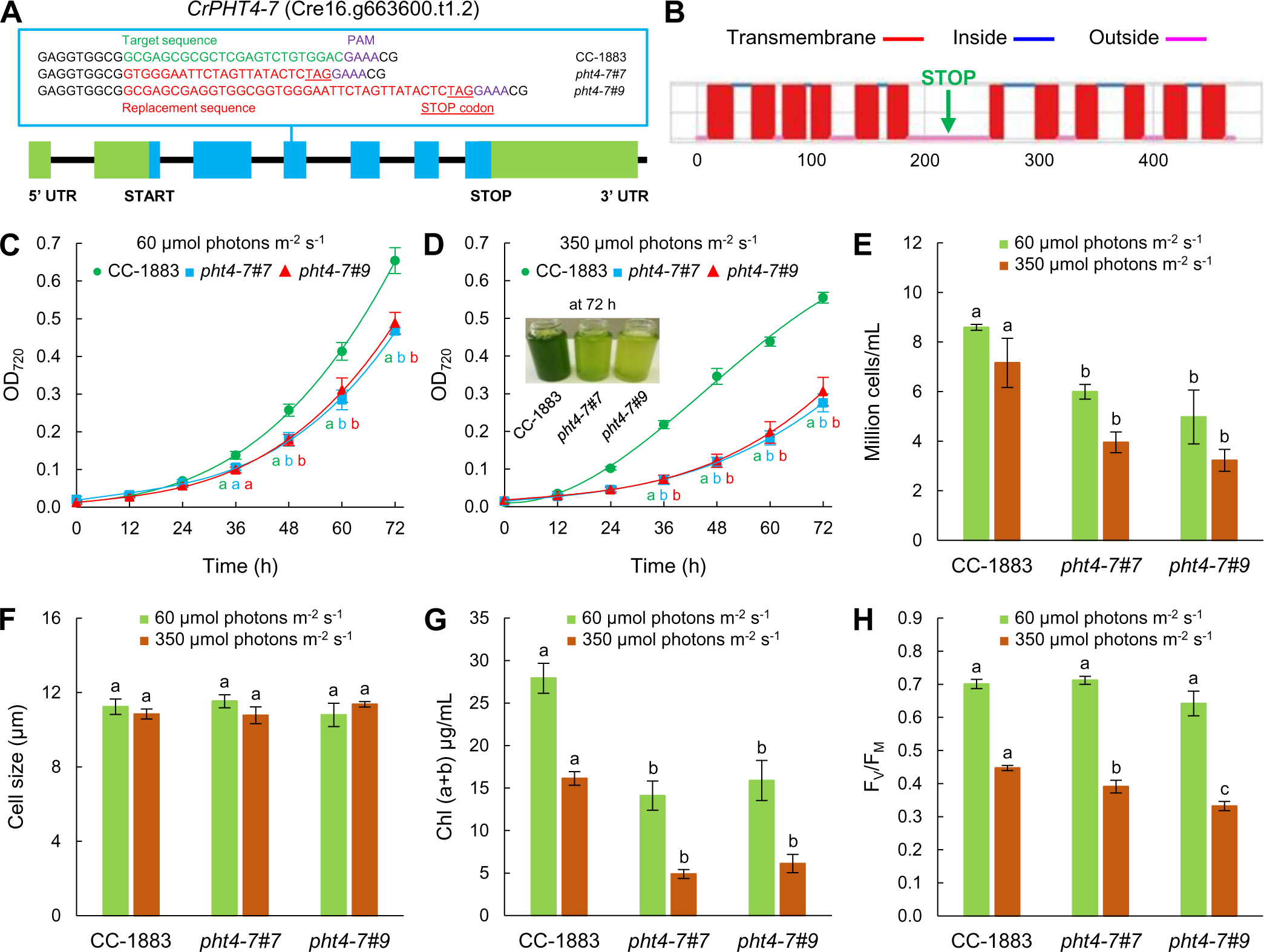
*pht4-7* mutants generated via the CRISPR/Cas12a technique exhibit diminished fitness. **A,** Physical map of *CrPHT4-7* (obtained from Phytozome, v. 13) with the replacement sequence including a stop codon, and a PAM sequence in the third exon in the *Crpht4-7#7 and #9* mutants. Exons are shown as blue boxes, introns as black lines, and promoter/5’ UTR and terminator sequences as green boxes. **B,** Prediction of transmembrane helices of CrPHT4-7 by Deep TMHMM v. 1.0.24. The introduction of the stop codon prevents the translation of at least six transmembrane helices. **C,** Culture growth of *pht4-7* mutants and the CC-1883 wild type, in TAP medium in continuous illumination of 60 µmol photons m^-2^ s^-1^ at 23°C, bubbled with air for 72 h in a Multi-Cultivator photobioreactor. The initial Chl content was set to 0.5 µg Chl(a+b)/mL. **D,** Culture growth in TAP medium under continuous illumination of 350 µmol photons m^-2^ s^-1^ at 23°C, bubbled with air for 72 h in a Multi-Cultivator photobioreactor. The initial Chl content was set to 0.5 µg Chl(a+b)/mL. A photograph of an aliquot of the cultures after 72 h of growth is shown in the inset. **E,** Cell numbers at 60 and 350 µmol photons m^-2^ s^-1^ after 72 h of growth. **F**, Cell sizes at 60 and 350 µmol photons m^-2^ s^-1^. **G,** Chl(a+b) contents after 72 h of growth at 60 and 350 µmol photons m^-2^ s^-1^ in a photobioreactor. **H,** F_V_/F_M_ values after 72 h of growth at 60 and 350 µmol photons m^-2^ s^-1^. The averages are based on three to five independent experiments with two to six biological replicates in each. The significance of differences between means were determined by ANOVA with Tukey post-hoc test. The means with different letters are significantly different (P < 0.05).

In agreement with the preliminary experiments, a significant difference in biomass accumulation (as assessed by OD_720_) between the WT and *pht4-7* mutant lines was found when grown at normal light (60 µmol photons m^-2^s^-1^, measured inside the culture tube; Fig. 2C). At high light (350 µmol photons m^-2^s^-1^), the fitness penalty associated with the absence of *pht4-7* became even more pronounced (Fig. 2D). Accordingly, the cell number and the Chl concentrations of the cultures (Chl(a+b)/ml) measured after three days of growth were significantly lower in the mutants than in the WT at both 60 and 350 µmol photons m^-2^ s^-1^ (Figs. 2E,G). We noted that the cell sizes of the mutants and the WT were very similar at normal and high light (Fig. 2F).

The F_V_/F_M_ value, an indicator of photosynthetic performance (Schansker et al., 2014, Sipka et al., 2021), was approximately 0.65 - 0.7 in all genotypes at normal light (Fig. 2H), which is typical for *C. reinhardtii* (e.g. Bonente et al., 2012, Santabarbara et al., 2019). At intense illumination, the F_V_/F_M_ value was about 0.45 in the WT, indicating downregulation of photosynthetic electron transport possibly involving photoinhibition. The reduction of photosynthetic efficiency was more enhanced in the *pht4-7* mutants than in the WT strain (Fig. 2H). From the above data, we conclude that CrPHT4-7 is required for cellular fitness, particularly under intense illumination.

We have also performed measurements on cultures grown in photoautotrophic conditions, in high salt (HS) medium, at normal light with CO_2_ supplementation. The *pht4-7* mutants were found to have mild growth phenotypes, thus photoautotrophic conditions did not enhance their stress sensitivity at moderate light intensity (Suppl. Fig. 4).

### Is CrPHT4-7 an ascorbate or a phosphate transporter?

Since CrPHT4-7 shows high amino acid sequence similarity to the AtPHT4;4 Asc transporter, we decided to assess Asc metabolism and function. This analysis, and the consecutive ones were carried out on alga cultures grown in TAP medium in Erlenmeyer flasks, enabling cultivating many more cultures in parallel than in the Multi-Cultivator instrument. By determining the cell number and the Chl content of the cultures after three days of growth in the Erlenmeyer flasks, we could confirm that the *pht4-7* mutant cultures grow more slowly than the WT especially at high light (Suppl. Fig. 5). In comparison with the Multi-Cultivator instrument, the difference between the mutants and WT was milder, indicating that shake-flask culturing was less stressful for the cells than growing in the Multi-Cultivator (see also Materials and Methods).

The cellular Asc concentration was about 0.8 mM in the CC-1883 strain when grown at 80 µmol photons m^-2^s^-1^ (Fig. 3A), that is in the same range as in other *C. reinhardtii* WT strains (Vidal-Meireles et al., 2017, 2020). In the *pht4-7 #7* mutant, the Asc concentration was about 1.1 mM and in the *pht4-7#9* line it was about 1.8 mM. At 500 µmol photons m^-2^s^-1^, the Asc content increased three-fold in the WT, whereas an about a ten-fold increase was observed in both *pht4-7* mutants, reaching approximately 15 mM Asc in the cell (Fig. 3A).

**Figure 3.**
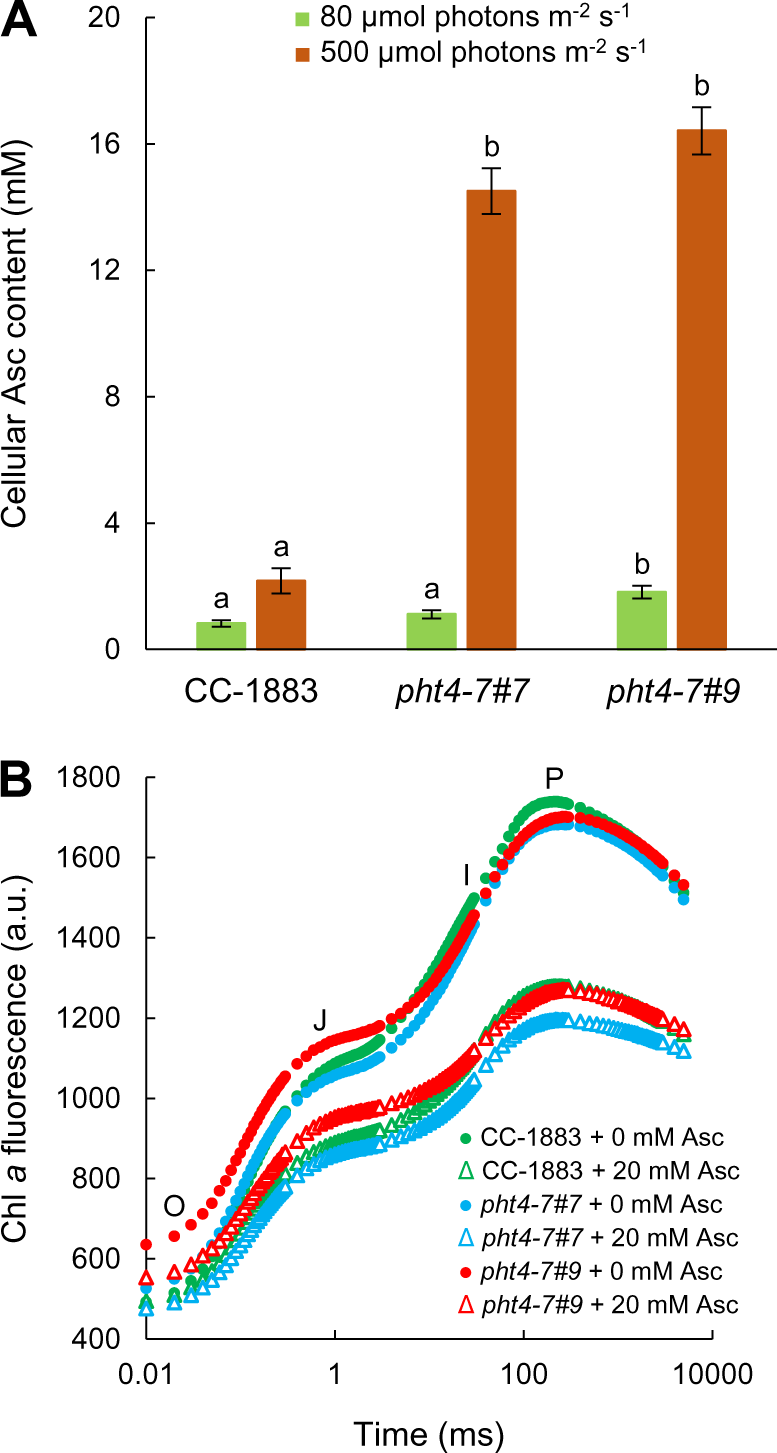
The *pht4-7* mutation leads to strong ascorbate (Asc) accumulation at high light and does not affect chloroplastic Asc uptake. **A,** Asc content of the *pht4-7* mutants and the CC-1883 strain after 72 h of growth in TAP medium at 80 and 500 µmol photons m^-2^ s^-1^. **B,** Fast Chl *a* fluorescence transients measured with or without 20 mM of Asc on cultures grown at 80 µmol photons m^-2^ s^-1^. The cultures were grown in Erlenmeyer flasks. The averages are based on three to six independent experiments with two to four biological replicates in each. The significance of differences between means were determined by ANOVA with Tukey post-hoc test. The means with different letters are significantly different (P < 0.05).

Next, we investigated the effect of Asc treatment on the fast Chl *a* fluorescence kinetics, which is a sensitive method to detect alterations in the function of the photosynthetic electron transport chain (e.g. Schansker et al., 2014). It was demonstrated earlier that a 10 mM Asc treatment causes a substantial, approx. 20-fold increase in cellular Asc content; at this high concentrations, Asc may inactivate the oxygen-evolving complex (OEC) in *C. reinhardtii* resulting in diminished variable Chl *a* fluorescence (Nagy et al., 2016; Nagy et al., 2018). We hypothesized that, if CrPHT4-7 is an Asc transporter in the chloroplast envelope membrane, then Asc transport into the chloroplast would be less efficient in its absence and consequently, less damage to the OEC should occur upon Asc treatment. As expected, the 20 mM Asc treatment resulted in a loss of variable fluorescence in cultures grown in normal light, but there were no clear differences between the WT and the *pht4-7* mutants (Fig. 3B). This result indicates that CrPHT4-7 does not contribute significantly to Asc transport into the chloroplast.

Ascorbate is a reductant for violaxanthin deepoxidase in vascular plants (Saga et al., 2010; Hallin et al., 2016), but is not required for green algal-type violaxanthin deepoxidases (Li et al., 2016; Vidal-Meireles et al., 2020). Instead, Asc mitigates an oxidative stress-related qI component of non-photochemical quenching (NPQ) and, therefore, NPQ is increased upon Asc-deficiency in *C. reinhardtii* (Vidal-Meireles et al., 2020). As expected, when the cultures were grown in normal light in TAP medium, the rapidly developing energy-dependent phase (qE) of NPQ was basically absent and NPQ mostly consisted of a slow phase, involving the zeaxanthin-dependent (qZ), state transition (qT) and the photoinhibitory (qI) components (e.g. Xue et al., 2015; Vidal-Meireles et al., 2020). The NPQ kinetics of the *pht4-7* mutants and the WT were similar in normal light (Fig. 4A). In high light, NPQ diminished remarkably in the *pht4-7* mutants relative to the WT (Fig. 4B). Since upon chloroplastic Asc-deficiency increased NPQ was observed due to the increase of qI (Vidal-Meireles et al., 2020), these results suggest that Asc transport into the chloroplast was maintained in the *pht4-7* mutants.

**Figure 4.**
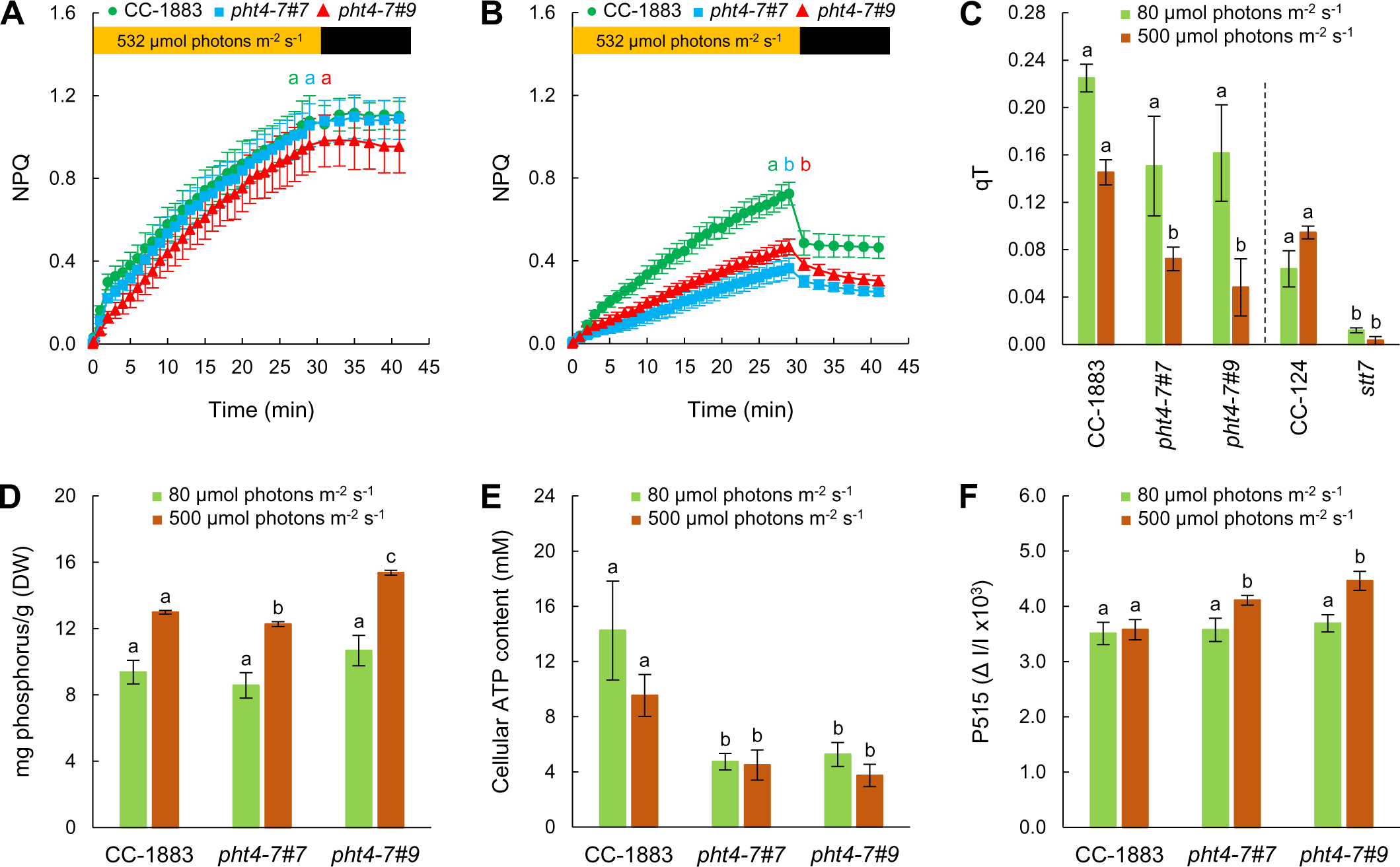
The *pht4-7* mutation alters photosynthetic redox homeostasis. **A,** NPQ of cultures grown in TAP medium at 80 µmol photons m^-2^ s^-1^. **B,** NPQ of cultures grown in TAP medium at 500 µmol photons m^-2^ s^-1^. For NPQ induction in panels A and B, light adaptation consisted of 30 min illumination at 532 µmol photons m^−2^s^−1^, followed by 12 min of dark adaptation interrupted with saturating pulses of 3000 µmol photons m^−2^s^−1^. **C,** State transition (qT, see the d in the Materials and methods section). **D,** Total phosphorous content. **E,** Cellular ATP content. **F,** Total proton motive force, determined based on the absorbance change at 515 nm against the 535 nm reference wavelength, expressed in ΔI/I units. All the cultures were grown in Erlenmeyer flasks. The averages are based on three to twelve independent experiments with one to two biological replicates in each. The significance of differences between means were determined by ANOVA with Tukey post-hoc test. The means with different letters are significantly different (P < 0.05). In the cases of panel A and B, significance was calculated at the end of the illumination period. In panel C, each mutant were compared to its own wild type. DW, dry weight.

We conducted state transition experiments using consecutive red and far-red illuminations in order to determine why NPQ was diminished in the *pht4-7* mutants (based on Ruban and Johnson, 2009; representative Chl *a* fluorescence traces can be found in Suppl. Fig. 6). The *pht4-7* mutants displayed reduced qT, especially under high light conditions (Fig. 4C), although to a lesser degree than a *stt7* state transition mutant (Fleischmann et al., 1999). This result raises the possibility that chloroplastic Pi may be decreased in the *pht4-7* mutants, since it has been described that state transition can be limited by Pi deficiency through insufficient LHCII phosphorylation (Petrou et al., 2008).

Next, measurements related to phosphate homeostasis were carried out. Phosphorous is taken up mostly in the form of Pi, therefore reduced Pi transport into the cell should decrease both the inorganic and organic phosphorous contents. We used ICP-OES to determine the total cellular phosphorous content and found that at normal light it was unaltered in the *pht4-7* mutants, whereas at high light, it was slightly diminished in the *pht4-7#7* mutant and augmented in the *pht4-7#9* mutant relative to the WT (Fig. 4D). Consequently, these data indicate that the absence of PHT4-7 did not limit phosphorous uptake into cells. On the other hand, inorganic phosphate is essential for ATP synthesis, and if CrPHT4-7 is a Pi transporter in the chloroplast envelope membrane, then its absence could limit ATP synthesis. Indeed, we found that cellular ATP content decreased in both *pht4-7* mutants, both in normal and high light conditions (Fig. 4E).

ATP production in the chloroplast is driven by transthylakoid proton motive force (pmf) that is physiologically stored as a ΔpH and a membrane potential (ΔΨ) (Cruz et al., 2005). Decreased chloroplastic phosphate availability, thereby ATP production (Carstensen et al., 2018) is expected to lead to increased pmf across the thylakoid membrane, especially in strong light (Cruz et al., 2005). As shown in Fig. 4F, total pmf is increased in both mutants at high light conditions, supporting this scenario.

As a next step, sensitivity to Pi limitation was assessed. Spot test, using four different Pi concentrations (0.2, 2, 100 and 200% of regular TAP medium) revealed that the growth of the *pht4-7* mutant strains was severely compromised upon Pi limitation in comparison with the WT (Fig. 5A). In liquid TAP cultures containing 0.5% Pi, cell proliferation was significantly diminished in the *pht4-7* mutants, as assessed by the Chl(a+b) content and cell number of the cultures (Fig. 5B,C).

**Figure 5.**
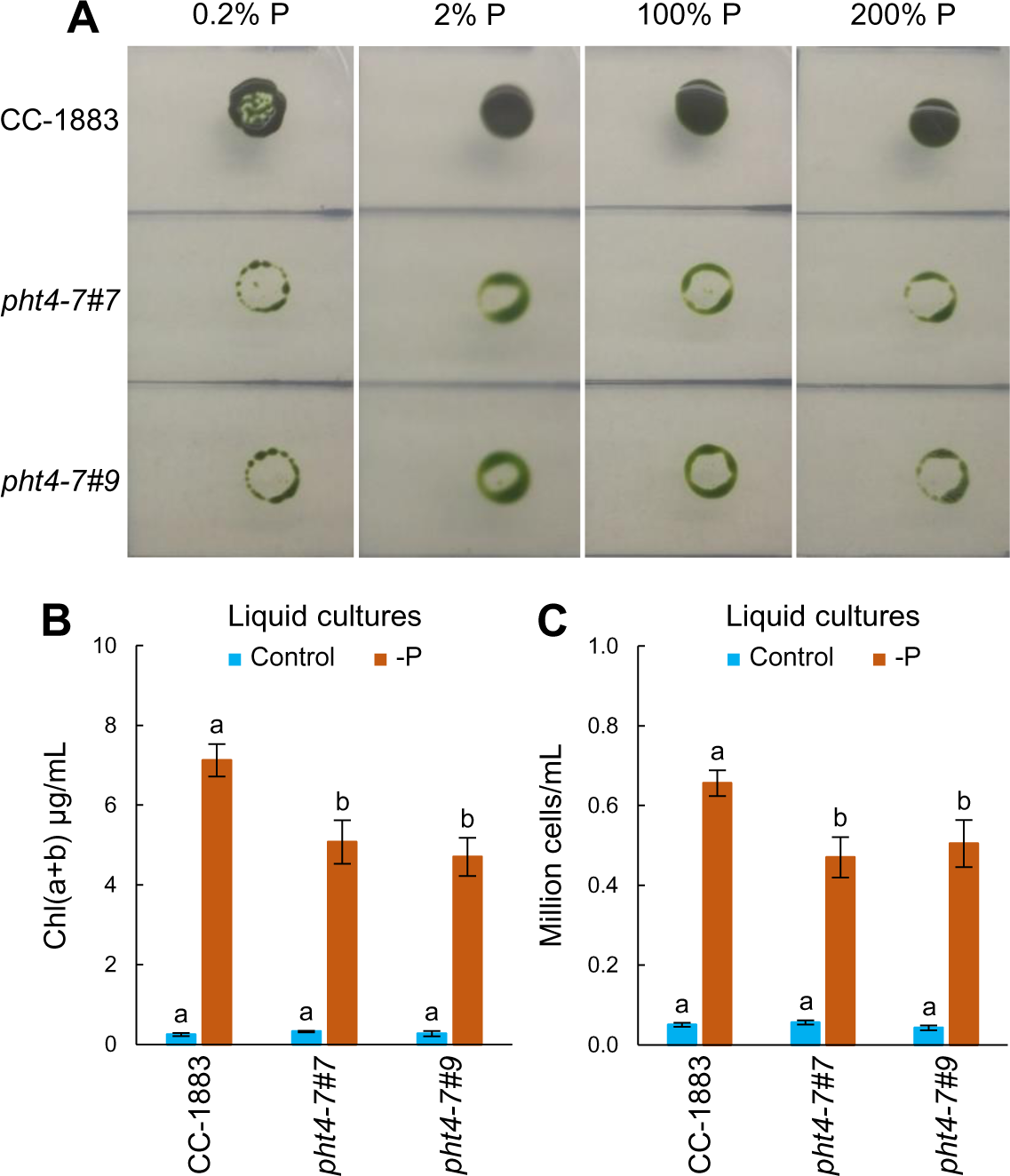
The *pht4-7* mutation leads to enhanced sensitivity to phosphorous limitation. **A,** Growth test of *pht4-7* mutants and the wild type strain on TAP agar plates containing different amounts of phosphorous; the photos were taken after 6 days. **B,** Chl(a+b) contents at the beginning and after 6 days phosphorous deprivation. **C,** Cell numbers at the beginning and after 6 days phosphorous deprivation. In panels B and C, liquid cultures were grown in Erlenmeyer flasks at 80 µmol photons m^-2^ s^-1^. The averages are based on five to ten independent experiments with one to two biological replicates in each. The significance of differences between means were determined by ANOVA with Tukey post-hoc test. The means with different letters are significantly different (P < 0.05).

Regarding the photosynthetic activity, we found that six days of Pi deprivation decreased the F_V_/F_M_ values very strongly (to about 0.1), and the decrease was slightly stronger in the mutants than in the WT (Fig. 6A). Importantly, the decrease in F_V_/F_M_ was caused by a very strong increase of the F_0_ value (Fig. 6B), indicating that the photosynthetic electron transport became reduced under Pi deprivation, most probably due to ATP deficiency (Fig. 4E), therefore a limited Calvin-Benson cycle activity. Upon the re-addition of Pi, the F_V_/F_M_ was almost fully restored within 24 h, showing that the downregulation of photosynthetic activity was reversible (Fig. 6A). Moreover, upon Pi limitation, NPQ increased, with the increase being less substantial in the *pht4-7* mutants than in the WT (Figs. 6C,D).

**Figure 6.**
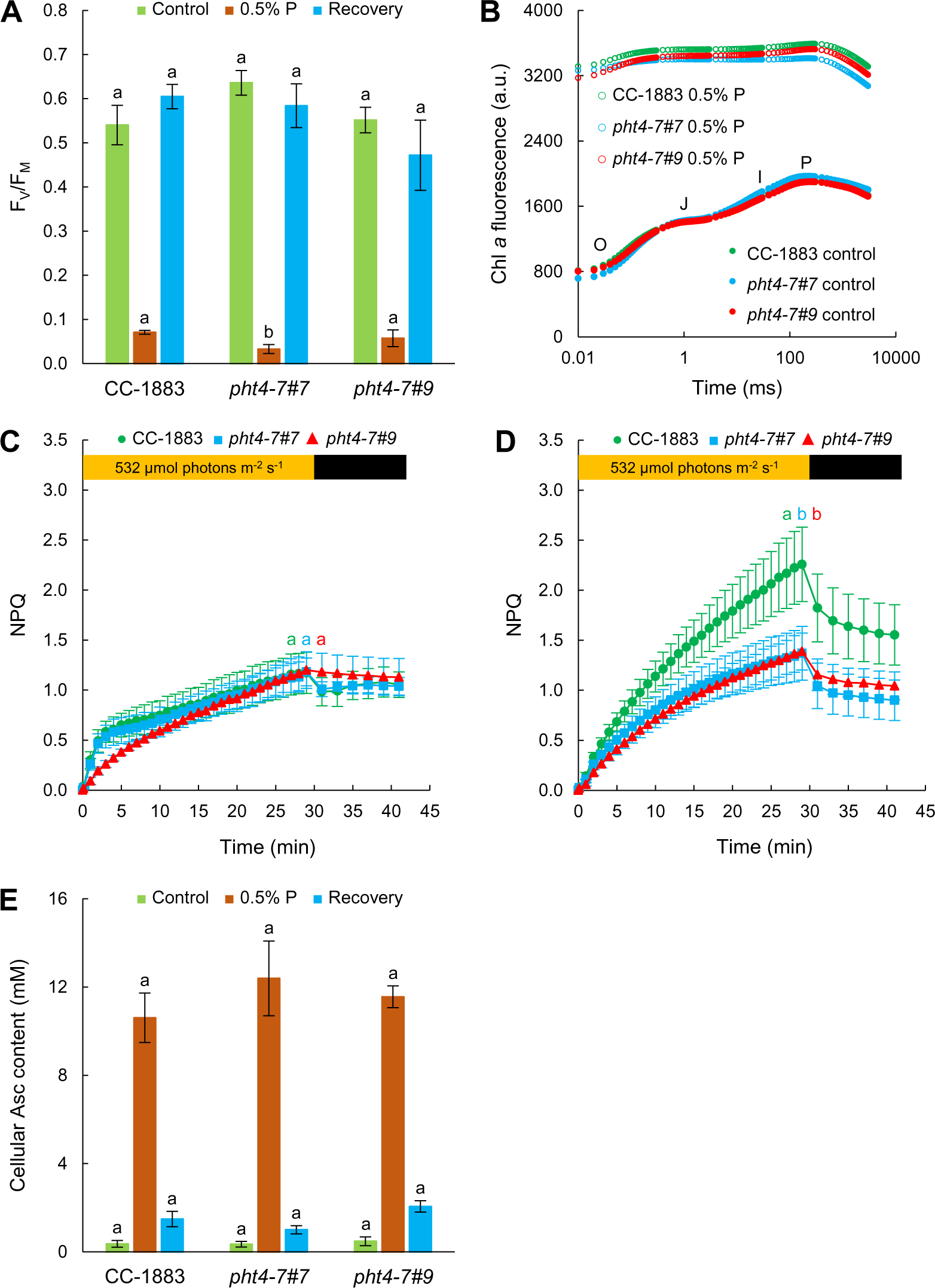
Alterations in photosynthetic activity upon phosphorous limitation. **A,** F_V_/F_M_ values of cultures grown in TAP and in TAP medium containing 0.5% P of regular TAP, for six days. For recovery, cultures were transferred to regular TAP media for one day. **B,** Fast Chl *a* fluorescence transients. **C,** NPQ (induced at 532 µmol photons m^−2^s^−1^) of cultures grown in regular TAP medium. **D,** NPQ of cultures grown in 0.5% P containing TAP medium for 6 days. **E,** Total cellular Asc contents. All the cultures were grown in Erlenmeyer flasks at 80 µmol photons m^-2^s^-1^. The same Chl(a+b) amounts were set for the Chl *a* fluorescence measurements. The averages are based on three to five independent experiments with one to two biological replicates in each. The significance of differences between means were determined by ANOVA with Tukey post-hoc test. The means with different letters are significantly different (P < 0.05). In the cases of panel A and B, significance was calculated at the end of the illumination period.

In addition, we have detected a very strong (about twenty to thirty-fold) increase in Asc contents upon Pi limitation in each strain, which was substantially restored by the re-addition of Pi within 24 h (Fig. 6E). These data show that Pi limitation leads to a strong Asc accumulation, similarly to sulphur deprivation involving oxidative stress (Nagy et al., 2018), and that upon the release of this stress effect, the Asc content rapidly returns to its original level.

### Genetic complementation and overexpression of CrPHT4-7

To confirm the relationship between the observed effects and the CrPHT4-7 mutation, genetic complementation experiments were carried out. We cloned the full-length CrPHT4-7 cDNA between the promoter and terminator sequence of *PSAD*, and subsequently transformed the *pht4-7* mutants with this construct (Fig. 7A). The complementation rescued the slow growth phenotype of the *pht4-7* mutants in at least 70% of the transformants tested (randomly selected lines for the complemented *pht4-7#7* mutant are shown in Suppl. Fig. 7A). The restored growth phenotype was also associated with higher Chl(a+b)/ml contents, improved photosynthetic performance (as assessed by the F_V_/F_M_ value), and moderate increases in Asc content, when grown in high light (Suppl. Figs. 7 B-D). Importantly, the complemented lines grew similarly upon Pi limitation in normal light as the WT (Suppl. Fig. 7E).

**Figure 7.**
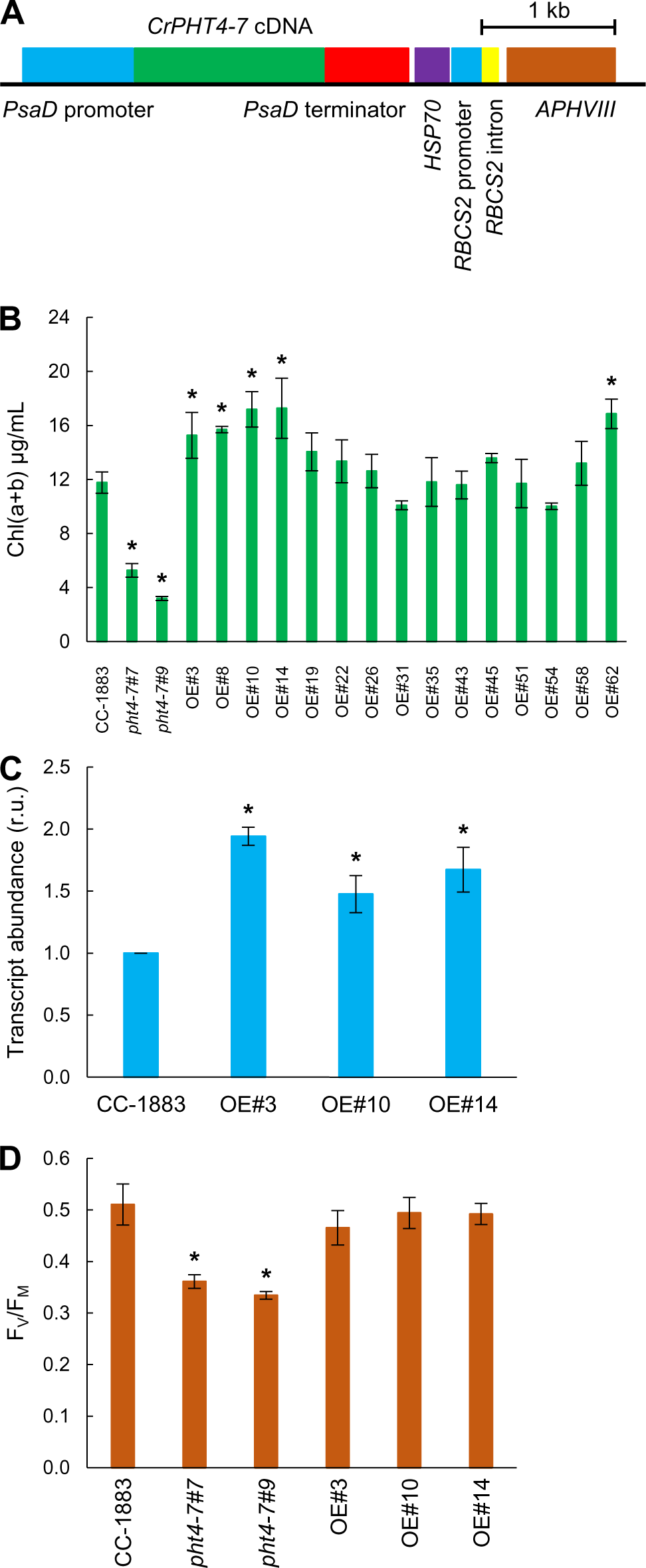
Overexpressing CrPHT4-7 in CC-1883 leads to improved growth in high light. **A,** Map of the pJR101 plasmid containing the coding sequence of *CrPHT4-7*, the strong *PSAD* promoter, the *APHVIII* resistance gene and the *PSAD* terminator. **B,** Chl(a+b) contents of CC-1883, *pht4-7* mutants, and several randomly selected *pht4-7*-overexpressing lines after three days of growth at 500 µmol photons m^-2^ s^-1^ in TAP medium in Erlenmeyer flasks. **C,** *PHT4-7* transcript abundance in CC-1883 and the selected *pht4-7*-overexpressing lines (OE#3, OE#10, OE#14) **D,** F_V_/F_M_ values measured on the same cultures. The averages are based on three to six independent experiments with two to six replicates in each. The significance of differences between means were determined by ANOVA with Dunette post-hoc test. Asterisks indicate significantly different means (p *<* 0.05) compared to the control strain CC-1883.

We also transformed CC-1883 with the above-mentioned construct to obtain CrPHT4-7-overexpressing lines. Out of 15 randomly selected lines, five showed significantly improved growth relative to the WT, as evidenced by higher Chl(a+b)/ml contents when grown in high light (Fig.7B). The relative transcript abundance of *PHT4-7* was significantly increased in the selected overexpressing lines (Fig. 7C). The F_V_/F_M_ values of the WT and the overexpressing lines did not differ significantly under high light treatment (Fig. 7D), indicating that the performance of the photosynthetic apparatus was similar in the overexpressing lines and the WT.

### Expression of CrPHT4-7 in a yeast strain lacking phosphate transporters

In order to study the substrate specificity of CrPHT4-7, we used the EY917 yeast strain in which five Pi transporters (PHO84, PHO87, PHO89, PHO90, PHO91) were inactivated, and the *GAL1* promoter drives the expression of *PHO84* enabling growth on galactose-containing media (Wykoff and O’Shea 2001). The EY917 strain lacking the five Pi transporters is considered conditional lethal, because spores are unable to germinate in the absence of galactose (i.e., on normal glucose-containing growth media). Plant phosphate transporters have been successfully investigated using Pi transporter-deficient yeast strains (Wang et al., 2015, Chang et al., 2019).

We transformed the EY57 (WT) and EY917 strains with the p426-TEF plasmid containing the *CrPHT4-7* gene (Fig. 8A). As controls, we used the EY57 and EY917 yeast strains transformed with the empty vector. The effect of expressing *CrPHT4-7* on the growth characteristics was then analyzed on glucose-containing medium. We found that the growth of the yeast strain expressing CrPHT4-7 was remarkably improved relative to the EY917 empty vector strain. Expressing CrPHT4-7 in the control strain EY57 had no significant effect on its growth properties in comparison with the EY57 empty vector strain (Fig. 8B). These data demonstrate that CrPHT4-7 acts as a Pi transporter.

**Figure 8.**
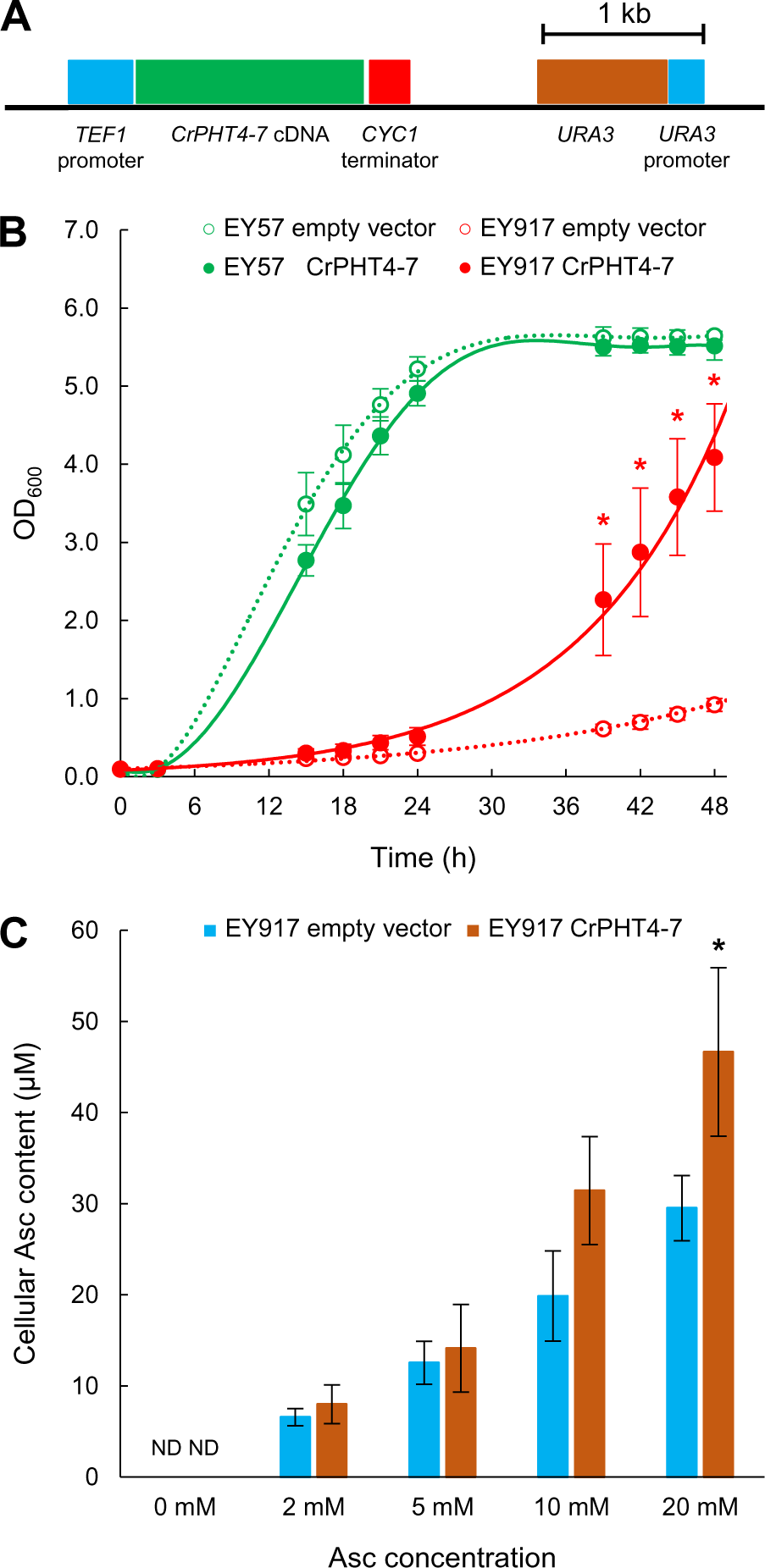
CrPHT4-7 transports phosphate in a yeast experimental system. **A,** Physical map of the construct for heterologous complementation. **B,** Growth rates of strain EY57 and the phosphate-transporter deficient strain EY917 expressing the empty vector or CrPHT4-7. **C,** Uptake of ascorbate (Asc) into yeast cells expressing CrPHT4-7 in comparison to the control strain. The cultures were incubated with 0, 2, 5, 10, 20 mM Asc for 15 minutes. The averages are based on three to four independent experiments. Data were analyzed by Welch’s unpaired *t*-test. Asterisks indicate significantly different means (p *<* 0.05) compared to the respective empty vector-containing strain. ND – non-detectable.

Since CrPHT4-7 is also a potential Asc transporter because it shares high similarity with the Asc transporter AtPHT4;4, its Asc uptake activity was also investigated in yeast cells. To this end, yeast cultures expressing CrPHT4-7 or an empty vector were incubated in the presence of 2, 5, 10 and 20 mM Na-Asc for 15 min. The control cultures contained no Asc, in agreement with published reports that yeast contains no Asc, but erythroascorbate instead (Spickett et al., 2000). At the two lowest concentration levels (2 and 5 mM), no significant difference between the EY917 and the CrPHT4-7 expressing yeast strains were observed. At 10 and 20 mM concentration levels, the uptake in the CrPHT4-7 expressing strain was more enhanced; however, the intracellular Asc concentration was only approx. 20 to 50 µM, i.e. approx. 0.2% of the external Asc level. This shows that Asc uptake by yeast cells is very moderate and it is only slightly increased by CrPHT4-7. We also note that the regular Asc content in *C. reinhardtii* in the range of 0.1 to 1 mM (Vidal-Meireles et al., 2017, and Fig. 3A), making it also unlikely that CrPHT4-7 substantially contributes to Asc content into the chloroplasts of *C. reinhardtii.* These data are in agreement with our results obtained with Chlamydomonas cells (Fig. 3B) and indicate that CrPHT4-7 does not act as an effective Asc transporter.

## Discussion

Phosphorus is an essential macronutrient fulfilling a wide range of functions for all living organisms, including microalgae. It is an essential compound of proteins, sugar phosphates, nucleic acids, and structural phospholipids and it is also required for information transfer via signal cascades (Dyhrman 2016). Phosphorous is of limited availability in nature; therefore, efficient P acquisition and storage, as well as the ability to cope with P limitation, are among the key factors determining the geographical distribution of single-celled phototrophs.

Cellular metabolism is severely affected by P starvation, including a slowdown of growth, changes in protein, lipid and starch biosynthesis and degradation, cellular respiration, and recycling of internal structures and compounds (reviewed by Sanz-Luque and Grossman, 2023). Ribosome degradation and a decrease in photosynthetic electron transport activity, including the loss of the PSII subunit, PsbA, has been also observed (Couso et al., 2020, Wykoff et al., 1998). This downregulation of photosynthetic electron transport helps to minimise photodamage, as due to the diminished Calvin-Benson cycle activity, much of the absorbed light energy cannot be used to support cell metabolism. Under P starvation, a number of photoprotective, P storage and uptake mechanisms are activated (reviewed by Sanz-Luque and Grossman, 2023).

Transporters play an essential role in Pi uptake and distribution within the cell. The Pi transporters situated in the cytoplasmic membrane of the cell are divided into two categories according to their affinity to the translocated Pi: there are low-rate high-affinity and high-rate low-affinity transporters, of which high-affinity Pi transporters are upregulated during P shortage (Grossman and Aksoy 2015). These include putative H^+^/PO_4_^3−^ PTA and Na^+^/PO_4_^3−^ PTB symporters (Moseley et al., 2006, Wang et al., 2020, Sanz-Luque and Grossman, 2023). In addition to PTA and PTB transporters, PHT3 and PHT4 transporters have also been identified by genetic analysis (e.g. Wang et al., 2020), but to our knowledge, none have been characterized in detail.

We found that CrPHT4-7, a member of the PHT4 family in *C. reinhardtii,* is a Pi transporter localized to the chloroplast envelope membrane (Fig. 1). The *pht4-7* mutants displayed retarded growth, compromised high-light tolerance, diminished ATP content and enhanced sensitivity towards Pi deprivation (Figs. 2, 4-6), demonstrating that CrPHT4-7 is required for maintaining Pi homeostasis in the chloroplast and for cellular fitness. These effects were particularly apparent under high light conditions. The reason could be that, at high light, Pi limitation within the chloroplast leads to relative ATP shortage (Fig. 4E), thereby limiting Calvin-Benson cycle activity and causing slower growth. At the same time, the photosynthetic electron transport chain becomes over-reduced, as indicated by the increased pmf (Fig. 4F). The decrease of the F_V_/F_M_ value in high light (Fig. 2H) indicates that in addition to the over-reduced electron transport chain, PSII may become also photoinhibited. Similar observations have been made upon Pi deficiency in green algae and in higher plants (Wykoff et al., 1998, Petrou et al., 2008; Carstensen et al., 2018) and in a *pht2;1* mutant of wheat (Guo et al., 2013).

Phosphate transporter mutants of vascular plants display enhanced NPQ due to a higher ΔpH induced by ATP limitation (Guo et al., 2013; Karlsson et al., 2015). By contrast, in our *pht4-7* mutants, NPQ decreased when grown in high light. NPQ mechanisms in green algae differ in many respects from those in vascular plants (Erickson et al., 2015, Vecchi et al., 2020). qE, which is a rapid ΔpH-dependent component appearing mostly under photoautotrophic growth conditions (Erickson et al., 2015), was not induced under our conditions (Fig. 4). Instead, NPQ developed on a timescale of several minutes, which may include the zeaxanthin-dependent (qZ), state transition-related (qT), and photoinhibitory (qI) components of NPQ (Erickson et al., 2015, Vidal-Meireles et al., 2020). We observed that pmf was elevated in both *pht4-7* mutants, suggesting that the decreased NPQ was not due to lack of membrane energization. On the other hand, ATP production and state transition (responsible for the qT component) were diminished in the *pht4-7* mutants (Fig. 4), most probably due to a limited Pi availability (as observed previously in *Dunaliella* upon Pi starvation, Petrou et al., 2008). Compromised state transition, acting as a major photoprotective mechanism in green algae (e.g., Goldschmidt-Clermont and Bassi 2015), may also explain the diminished F_V_/F_M_ values in the *pht4-7* mutants grown at high light (Fig. 2).

The apparent Pi limitation in the chloroplast led to a dramatic increase in cellular Asc content when the cultures were grown in high light (Fig. 3A). The high-level accumulation of Asc in the *pht4-7* mutants may occur to mitigate reactive oxygen species, as provoked by compromised state transition and ATP synthesis diminishing CO_2_ fixation. When accumulating to high levels, Asc may also inactivate the OEC to alleviate the consequences of over-reduction of the electron transport chain when CO_2_ assimilation is impaired (Nagy et al., 2018). Thus, it seems that chloroplastic Pi-deficiency triggers high Asc accumulation in *C. reinhardtii*, similar to induction of Asc accumulation upon sulfur deprivation (Nagy et al., 2016). Conversely, overexpression of CrPHT4-7 in *C. reinhardtii* resulted in enhanced resistance to high light stress, demonstrating that Pi transport can limit photosynthesis under intensive illumination.

Although CrPHT4-7 exhibits a relatively high degree of similarity with AtPHT4;4, it did not show significant Asc transport activity. In algal cells, Asc uptake into the chloroplasts, as tested by incubating the cultures with Asc and measuring Chl *a* fluorescence transients, did not seem to differ between the WT and the *pht4-7* mutants. When expressed in yeast, CrPHT4-7 did not enhance Asc uptake into the cells in the physiologically relevant concentration range (Figs. 3, 8). At high concentrations, there was a slight enhancement of Asc uptake by the CrPHT4-7 transporter; however, physiologically, it is probably of little significance.

In summary, we have shown that CrPHT4-7 supports Pi homeostasis and photosynthesis in the chloroplasts and overexpressing CrPHT4-7 enhanced high light tolerance. On the other hand, the loss of CrPHT4-7 function was not lethal even though Pi is essential to maintain chloroplast function. It thus appears likely that there are additional PHT transporters located in the chloroplast envelope membrane. PHT2 transporters are not found in green algae (Bonnot et al., 2017), therefore, other members of the PHT4 family are likely to supply Pi to chloroplasts, as suggested also by *in silico* analysis (Wang et al., 2020, Wang et al., 2023). Confirming the identity and revealing the physiological roles of additional chloroplastic Pi transporters should be the subject of future studies. Pi transporters located in the plasma membrane, the mitochondria, and other cellular compartments have not been characterized in detail in green algae; their analysis will be important to fully exploit the so-called “luxury uptake” characteristics of green algae towards mitigating excess Pi in polluted waters and for the development of new wastewater treatment strategies.

## Materials and Methods

### Algal strains

The *pht4-7#7* and *pht4-7#9* mutant strains had been generated via CRISPR/Cas12a, published previously, using CC-1883 as the background strain (Ferenczi et al., 2017). To generate complementation and PHT4-7 overexpressing lines, the coding sequence of the *CrPHT4-7* gene was synthesized (GeneCust, Boynes, France) with NdeI and EcoRI restriction sites at the 5’ and 3’ ends, respectively. The fragment was cloned into the similarly digested vector pJR39 (Neupert et al., 2009), generating the transformation vector pJR101. Nuclear transformation of the CC-1883 and *pht4-7#7* strains of *C. reinhardtii* was performed using the glass bead method (Neupert et al., 2012). Selection was performed on TAP plates supplemented with 10 µg/mL paromomycin.

### Generation of PHT4-7 expressing yeast strains

We used the EY57 (*MATa ade2-1 trp1-1 can1-100 leu2-3,112 his3-11,15 ura3*) and the EY917 (*MATα ade2-1 trp1-1 can1-100 leu2-3,112 his3-11,15 ura3 pho84::HIS3 pho87::CgHIS3 pho89::CgHIS3 pho90::CgHIS3 pho91::ADE2, pGAL1-PHO84* (EB1280)) *S. cerevisiae* strains that were kindly provided by Dr. Dennis Wykoff (Villanova University, USA).

The coding sequence of the *CrPHT4-7* gene with BamHI and EcoRI restriction sites at the 5’ and 3’ ends was cloned into the similarly digested vector p426-TEF (containing *URA3* marker), generating the transformation plasmid. We transformed EY57 and EY917 *S. cerevisiae* strains with the plasmid containing the *CrPHT4-7* gene by selecting for the *URA3* marker. We followed the transformation protocol by Gietz and Schiestl (2007). For transformation, strains were grown in synthetic media lacking uracil and containing 2% galactose.

### Structure prediction of PHT4-7 and sequence alignment

To predict the transmembrane helices of CrPHT4-7, we used the TMHMM v. 2.0 (Krogh et al., 2001), Deep TMHMM v. 1.0.24 (Hallgren at al., 2022) and the Phyre^2^ v. 2.0 (Kelley et al., 2015) online softwares. Amino acid sequence alignment was performed by MultAlin (Corpet, 1988).

### Growth of alga cultures

Precultures were grown mixotrophically in Tris-acetate-phosphate medium (TAP, Gorman and Levine, 1965) in 25-mL Erlenmeyer flasks for three days on a rotatory shaker at 130 rpm, at 23°C and 80 µmol photons m^-2^ s^-1^, measured at the top of the flasks. By the third day of growth in TAP, a cell density of 2-4 million cells/mL was reached.

For the assessment of culture growth parameters (in Fig. 2), the precultures were diluted to 0.5 µg Chl(a+b)/mL and were placed in a Multi-Cultivator MC 1000-OD instrument (Photon Systems Instruments, Brno, Czech Republic). The cultures were grown for up to three days at 23°C with intense air bubbling, at a light intensity of 60 or 350 µmol photons m^-2^ s^-1^ measured within the culture tubes.

For measuring the rest of the physiological measurements (e.g. photosynthetic parameters, ATP and Asc contents), the cultures were grown in 50-mL Erlenmeyer flask for three days on a rotatory shaker at 130 rpm, 23°C. For most experiments, the cultures were grown in TAP medium, and in a subset of experiments high salt (HS) medium was used. The initial Chl concentration was 0.5 µg Chl(a + b)/mL, and the light intensity was 80 or 500 µmol photons m^-2^ s^-1^, measured at the top of the flasks (the effective light intensity is remarkably lower within the flask). We noted that shake-flask culturing was less stressful for the cells than growth in the Multi-Cultivator MC 1000-OD instrument.

### Growth of yeast cultures for CrPHT4-7 expression

In order to enable the growth of the EY917 strain (containing *GAL1-PHO84*), precultures for both strains (EY57 and EY917) were grown in synthetic yeast media with 2% galactose and appropriate amino acids for one day on a rotatory shaker at 30°C. To prevent *PHO84* expression, the precultures were harvested by centrifugation (3000 g, 1 min, 25°C), washed two times, and were diluted to OD_600_ = 0.1 with synthetic yeast media containing 2% glucose and appropriate amino acids without uracil. The cultures were grown for two days on a rotatory shaker at 140 rpm at 30°C.

### Chlorophyll and Asc content measurements and phosphorus content determination in C. reinhardtii

Chl(a+b) content was determined according to Porra et al. (1989), and the Asc content was determined as in Kovács et al., (2016). Total phosphorus content determination was performed by ICP-OES, as described in Nagy et al. (2018).

### ATP content determination

ATP was measured using the Adenosine 5’-triphosphate (ATP) Bioluminescent Assay Kit (Sigma-Aldrich) according to the instructions of the manufacturer. 3×10^7^ algal cells were harvested by centrifugation (21130 g, 1 min, 4°C) and washed once with ice cold sterile water The pellets were resuspended in 250 µl ice cold sterile water. Cells were broken by vortexing for 2 min with 80 µl quartz sand. After the vortexing the samples were centrifuged (21130 g, 1 min, 4°C). 200 µl of the supernatant were transferred into EZ-10 Spin Columns (Bio Basic Inc.) and rapidly spun down (21130 g, 1 min, 4°C). Until ATP determination the samples were stored on ice. The cellular ATP concentration was determined using a cell volume of 140 femtoliters (Craigie and Cavalier-Smith, 1982).

### Phosphorus deprivation

Precultures were grown mixotrophically in TAP medium in 50 mL Erlenmeyer flasks for three days on a rotatory shaker at 130 rpm, 23°C and 80 µmol photons m^-2^ s^-1^. After three days the cells were harvested by centrifugation (3000 g, 1 min, 23°C), washed three times, and were diluted to 0.5 µg/mL Chl (a+b) with 0.5% Pi-containing TAP medium. The cultures were grown at 23°C, 80 µmol photons m^-2^ s^-1^, on a rotatory shaker at 130 rpm, for six days.

### Drop test

The growth characteristics of the strains were tested on TAP agar plates, containing different amounts of phosphorus (2,04 µM - 0,2%; 20,4 µM - 2%; 1,02 mM - 100% 2,04 mM - 200%). Precultures were grown mixotrophically in TAP medium in 50 mL Erlenmeyer flasks for three days on a rotatory shaker at 130 rpm, 23°C and 80 µmol photons m^-2^ s^-1^. After three days the cells were harvested by centrifugation (3000 g, 1 min, 23°C), washed three times, and were diluted to 5 µg/mL Chl (a+b) with 0.5% Pi-containing TAP medium. 10 µL of each algal strain was dropped onto the agar plates. The plates were incubated at 23 °C for 6 days. The intensity of illumination was 80 µmol photons m^-2^ s^-1^.

### Ascorbate uptake measurements in C. reinhardtii and yeast

The three days old *C. reinhardtii* precultures (in TAP medium) were diluted to 10 µg/mL Chl (a+b), and incubated for two hours on a rotatory shaker at 23°C and 80 µmol photons m^-2^ s^-1^ with or without 20 mM Asc.

Yeast cultures were kept in yeast synthetic media with 2% glucose and appropriate amino acids for one day on a rotatory shaker at 30 °C. After one day we measured the OD_600_ values of the cultures (the strains were grown to log phase OD_600_ = 0.7 - 1.5), and set OD_600_ = 0.8. We added 0, 2, 5, 10, 20 mM Asc, and incubated the cultures for 15 minutes on a rotatory shaker at 30°C. We harvested the cells by centrifugation (3000 g, 1 min, 4°C), washed three times with 40 mL ice cold synthetic media, and immediately frozen in liquid nitrogen. Cells were broken by vortexing for 30 s with glass beads (425-600 µm, Sigma-Aldrich, St. Louis, USA). The Asc content was determined as in Kovács et al., (2016) with slight modifications.

### Analysis of gene expression

For isolation of RNA, 2 ml of cultures were harvested and Direct-Zol RNA MiniPrep kit (Zymo Research) was used, following the recommendations of the manufacturer. To remove contaminating DNA from the samples, the isolated RNA was treated with DNaseI (Zymo Research). RNA integrity was checked on a 1% (w/v) denaturing agarose gel. 1 µg of total RNA was used for cDNA synthesis with random hexamers using FIRESript reverse transcriptase (Solis BioDyne). To confirm the absence of DNA contaminations, an aliquot of the RNA sample was used without reverse transcriptase. Real-time qPCR analysis was performed using a Bio-Rad CFX384 Touch Real-Time PCR Detection System, using HOT FIREPol EvaGreen qPCR Mix Plus (Solis Biodyne) for cDNA detection. The primer pairs for the reference genes (*actin* [Cre13.g603700], *β-Tub2* [Cre12.g549550], *CBLP* [Cre06.g278222], *UBQ2* [Cre09.g396400]) used in RT-qPCR were published earlier in Vidal-Meireles et al. (2017). For *PHT4-7* 5’-CAACTGGGGCTACTACACGC-3’ forward and 5’-CCATGACCCGCTCCTCATATC-3’ revers primers were used. The data are presented as fold-change in mRNA transcript abundance, normalized to the average of the reference genes, and relative to the WT sample. Real-time qPCR analysis was carried out with three technical replicates for each sample and three to four biological replicates were analysed. The standard errors (SE) were calculated based on the different transcript abundances amongst the independent biological replicates.

### Determination of cell size and cell number

The cell size and cell number were determined by a Luna-FL™ dual fluorescence cell counter (Logos Biosystems Inc.).

### Chl a fluorescence measurements

Fast chl *a* fluorescence measurements were carried out with a Handy-PEA instrument (Hansatech Instruments Ltd, King’s Lynn, UK), as described in Nagy et al. (2018).

Non-photochemical quenching was measured using a Dual-PAM-100 instrument (Heinz Walz GmbH). *C. reinhardtii* cultures were dark adapted for 30 min on a rotatory shaker; then, liquid culture containing 40 µg Chl(*a*+*b*)/mL was filtered onto Whatman glass microfiber filters (GF/B) that were placed between two microscopy coverslips with a spacer to allow for gas exchange. For NPQ induction, light adaptation consisted of 30 min illumination at 532 µmol photons m^−2^s^−1^, followed by 12 min of dark adaptation interrupted with saturating pulses of 3000 µmol photons m−^2^s−^1^.

For analyzing state transition, actinic red light (AL, 15 μmol photons m^-2^ s^-1^) and far red (FR) light (255 μmol photons m^-2^ s^-1^) were employed for 15 min (phase 1) on dark-adapted cultures. After this phase, the far red light was turned off and only red light illumination was employed for 15 min to induce state II (phase 2). Finally, we used again the red light - far red light combination for 15 min to drive the state II - state I transition (phase 3). During the measurement, saturating light pulses (8000 μmol photons m^-2^ s^-1^ for 600 ms) were given every minute. qT parameter was calculated as: qT = (F_M_ ^I^ - F_M_ ^II^)/F_M_ ^II^, in which F_M_ ^I^ was determined at the end of the phase 3, and F ^II^ at the end of the phase 2.

### Pmf measurements

Estimation of the trans-thylakoid proton motive force (pmf) was carried out by Dual-PAM-100 system with the P515/535 extended emitter-detector modules (Schreiber and Klughammer, 2008). Before the measurement, samples were kept for 10 min in darkness, and cultures equivalent to 40 µg/mL Chl (a+b) were filtered onto a GF/C filter paper. Samples were placed between two object slides with a spacer to allow for gas exchange. Samples were illuminated with 190 µmol photons m^-2^s^-1^ actinic red light for two minutes, then actinic light was switched off. The absorbance change at 515 nm against the 535 nm reference wavelength was recorded during the light-dark transition (Cruz et al., 2001; Kramer and Sacksteder, 1998). The change of signal was expressed in ΔI/I units (Schreiber and Klughammer, 2008).

### Generation of PHT4-7-Venus expressing lines and localization of PHT4-7 in Chlamydomonas

Nuclear transformation of strains UVM11 and CC-4533 (also known as cMJ030) of *C. reinhardtii* with the plasmid pLM005-CrPHT4-7 was done using the glass bead method (Neupert et. al., 2012). We also transformed the CC-1883 strain and the *pht4-7* mutants with this construct, but failed to obtain transgenic clones showing a clear Venus signal, most probably due to very low expression levels caused by epigenetic transgene silencing (Neupert et al., 2020).

The pLM005-CrPHT4-7 plasmid contains the full length CrPHT4-7 gene including the introns. The plasmid was linearized using the restriction enzyme ScaI. Pre-cultures of the transformed strains were grown mixotrophically in TAP medium in 25-mL Erlenmeyer flasks for three days. The strains were then transferred to Tris-phosphate (TP) medium and further grown for 16 hrs under the above-mentioned conditions, after which the cells were immobilized in 0.8% low-melt agarose (Carl Roth, Karlsruhe, Germany) before imaging. Imaging was performed using a Leica TCS SP8 confocal laser scanning microscope with a hybrid detector (Leica, Heidelberg, Germany). Single optical sections were taken using HCPLAPO CS2 63× (NA:1.2) water immersion objective with a working distance of 0.3 mm. Microscope configuration was as follows: scan speed: 200; line averaging: 4; scanning mode: unidirectional; zoom: 7×; excitation: 514 nm (Venus-CrPHT4-7), 552 nm (Chl auto-fluorescence). Venus-CrPHT4-7 fluorescence and Chl auto-fluorescence were detected between 520-540 nm and 650-750 nm respectively. HyD SP GaAsP detector was used to detect the Venus-CrPHT4-7 signal. Images were pseudocolored and analyzed using Leica LAS AF software (version 2.6) and ImageJ (version 1.53k).

### Statistics

The presented data are based on at least three independent experiments. When applicable, averages and standard errors (±SE) were calculated. Statistical significance was determined using Welch’s unpaired t-test (GraphPad Prism v. 10.0.2.232 online software), ANOVA with Tukey post-hoc test (OriginPro 2020b software) or Dunette post-hoc test (IBM SPSS Statistics v. 25.0 software). Changes were considered statistically significant at P < 0.05.

### Accession Numbers

The accession number for *C. reinhardtii PHT4-7* (also called *PHT7*) gene is Cre16.g663600.

## Acknowledgements

The authors thank Drs. Péter Horváth and Balázs Papp (BRC Szeged, Hungary) for laboratory equipment support and Dr. Cornelia Spetea (University of Gothenburg, Sweden) for the fruitful discussions. The authors also thank Miklós Prodán (TTK, Budapest, Hungary) for the assistance with phosphorus content determination and Dr. Dennis Wykoff (Villanova University, USA) for providing us with the yeast strains.

## Data Availability Statement

All data presented in this study are available within this article or Supplementary Materials. There are no special databases associated with this manuscript.

## Supporting Information

Additional Supporting Information may be found online in the Supporting Information section at the end of the article.

## Author contributions

SZT conceived the study with the contributions of AM and MCJ. DT, SK, AF, AVM, LK, LW, ZK, RT, EM, KS, and JN performed the experiments and data analysis. SZT wrote the manuscript with the contributions of DT, SK, JN, RB, MCJ, and AM.

All authors reviewed the manuscript and approved the final version.

## Funding

This work was supported by the Lendület/Momentum Programme of the Hungarian Academy of Sciences (LP2014/19 research grant to S.Z.T.) and the National Research, Development, and Innovation Office (K132600 research grant to S.Z.T.) A.F. was supported by Biotechnology and Biological Sciences Research Council (BBSRC) grant BB/R506163/1. L.W and M.J. were supported by U.S. Department of Energy Grant DE-SC0020195. M.J. is an Investigator of the Howard Hughes Medical Institute. J.N. and R.B. were supported by the Max Planck Society.

## Supplementary materials

**Suppl. Figure 1. Amino acid sequence alignment of members of the PHT4 family in *Arabidopsis thaliana* (AtPHT4) and CrPHT4-7 in *C. reinhardtii.*** Conserved amino acids are indicated in red. Predicted transmembrane regions are shown in green boxes. The amino acid sequence alignment and prediction of transmembrane helices were performed using the MultAlin and the Phyre2 v. 2.0 online software, respectively.

**Suppl. Figure 2. Subcellular localization of CrPHT4-7 in the CC-4533 strain. A,** Representative fluorescence microscopy images of CC-4533 and B, CC-4533 expressing pLM005-CrPHT4-7. Venus fluorescence and Chl autofluorescence were detected between 520-540 nm and 650-750 nm, respectively. The merged Venus + Chl autofluorescence image is also shown. Scale bar: 5 μm.

**Suppl. Figure 3. Culture growth of independent *pht4-7* mutant lines generated by the CRISPR/Cas12a technique in TAP medium in continuous illumination in a Multi-Cultivator photobioreactor. A**, Culture growth at 60 µmol photons m^-2^ s^-1^ as assessed by measuring optical density (OD) at 720 nm. B, Culture growth at 350 µmol photons m^-2^ s^-1^. The initial Chl content was set to 0.5 µg Chl(a+b)/mL, the temperature was kept at 23°C, and the cultures were bubbled with air.

**Suppl. Figure 4. Phenotype of *pht4-7* mutants under photoautotrophic growth conditions. A**, Culture growth of *pht4-7* mutants and the CC-1883 wild type, in HS medium in continuous illumination of 60 µmol photons m^-2^ s^-1^ at 23°C, bubbled with air for 72 h in a Multi-Cultivator photobioreactor. The initial Chl content was set to 0.5 µg Chl(a+b)/mL. B, F_V_/F_M_ values after 72 h of growth in HS medium at 60 µmol photons m^-2^ s^-1^. C, NPQ of cultures grown in HS medium at 60 µmol photons m^-2^ s^-1^. The averages are based on three independent experiments with one to two biological replicates in each. The significance of differences between means were determined by ANOVA with Tukey post-hoc test. The means with different letters are significantly different (P < 0.05). In the case of panel A significance was calculated for the last time point (72 h). In the case of panel C significance was calculated at the end of the illumination period.

**Suppl. Figure 5. Cell number and chlorophyll values of *pht4-7* mutants and the wild type grown in Erlenmeyer flasks. A,** Cell numbers after 72 h of growth at 80 and 500 µmol photons m^-2^ s^-1^. B, Chl(a+b) contents after 72 h of growth at 80 and 500 µmol photons m^-2^ s^-1^. C, µg Chl(a+b)/million cells values after 72 h of growth at 80 and 500 µmol photons m^-2^ s^-1^. The averages are based on 15 to 20 independent experiments with one to three biological replicates in each. The significance of differences between means were determined by ANOVA with Tukey post-hoc test. The means with different letters are significantly different (P < 0.05).

**Suppl. Figure 6. Typical state transition kinetics of *pht4-7* and *stt7* mutants. A-E,** Cultures were grown in TAP medium in Erlenmeyer flasks under continuous illumination of 80 µmol photons m^-2^s^-1^. F-J, Cultures were grown in TAP medium under continuous illumination of 500 µmol photons m^-2^s^-1^. F_M_ ^II^ and F ^I^ values were used to calculate qT (see Materials and Methods).

**Suppl. Figure 7. Complementation of the *pht4-7#7* CRISPR/Cas12a mutant. A,** Phenotype of the CC-1883 strain, the *pht4-7#7* mutant, and several randomly selected complementation lines grown for three days at 500 µmol photons m^-2^ s^-1^. B, Chl(a+b) contents of the CC-1883 strain, the *pht4-7#7* mutant, and two selected complementation lines (*C#23*, *C#24*) at 500 µmol photons m^-2^ s^-1^. C, F_V_/F_M_ values under the same conditions. D, Ascorbate accumulation under the same conditions. E, Chl(a+b) contents after six days of phosphate deprivation at 80 µmol photons m^-2^ s^-1^. The cultures were grown in Erlenmeyer flasks. The averages are based on four to ten independent experiments. The significance of differences between means were determined by ANOVA with Tukey post-hoc test. The means with different letters are significantly different (P < 0.05).

